# Plasticity in auditory categorization is supported by differential engagement of the auditory-linguistic network

**DOI:** 10.1101/663799

**Authors:** Gavin M. Bidelman, Breya Walker

## Abstract

To construct our perceptual world, the brain categorizes variable sensory cues into behaviorally-relevant groupings. Categorical representations are apparent within a distributed fronto-temporo-parietal brain network but how this neural circuitry is shaped by experience remains undefined. Here, we asked whether speech (and music) categories might be formed within different auditory-linguistic brain regions depending on listeners’ auditory expertise. We recorded EEG in highly skilled (musicians) vs. novice (nonmusicians) perceivers as they rapidly categorized speech and musical sounds. Musicians showed perceptual enhancements across domains, yet source EEG data revealed a double dissociation in the neurobiological mechanisms supporting categorization between groups. Whereas musicians coded categories in primary auditory cortex (PAC), nonmusicians recruited non-auditory regions (e.g., inferior frontal gyrus, IFG) to generate category-level information. Functional connectivity confirmed nonmusicians’ increased left IFG involvement reflects stronger routing of signal from PAC directed to IFG, presumably because sensory coding is insufficient to construct categories in less experienced listeners. Our findings establish auditory experience modulates specific engagement and inter-regional communication in the auditory-linguistic network supporting CP. Whereas early canonical PAC representations are sufficient to generate categories in highly trained ears, less experienced perceivers broadcast information downstream to higher-order linguistic brain areas (IFG) to construct abstract sound labels.

## INTRODUCTION

Mapping sensory cues in the environment onto common perceptual identities is a prerequisite for complex auditory processing. In speech perception, acoustically distinct sounds along a continuum of similar features are identified categorically in that they are heard as belonging to one of only a few discrete phonetic classes (Pisoni and Luce, 1987). Because categories represent knowledge of stimulus groupings, patterns, and the linkage of sensory cues with internal memory representations (Seger and Miller, 2010), it is argued that categorization reflects the nexus between perception and cognition (Freedman et al., 2001). Understanding how the brain imposes these “top-down” transformation(s) onto the “bottom-up” sensory input to construct meaningful categories is among the many broad and widespread interests to understand how sensory features are realized as invariant perceptual objects (Phillips, 2001; Pisoni and Luce, 1987).

The process of categorization requires a higher-order abstraction of the sensory input and consequently, offers an ideal window into how experiential factors might alter this fundamental mode of auditory perception-cognition. Both behavioral and neuroimaging studies demonstrate category-level sensitivity is malleable to listening experience, learning, and stimulus familiarity (Myers, 2014). While categorical perception (CP) emerges early in life (Eimas et al., 1971), it can be further modified by native language experience (Bidelman and Lee, 2015; Kuhl et al., 1992; Xu et al., 2006). Similarly, trained musicians show sharper categorical boundaries than their nonmusician peers for pitch intervals of the musical scale (Burns and Campbell, 1994; Burns and Ward, 1978; Zatorre, 1983). Conceivably, long-term experience (and increased familiarity) with the sounds of a certain domain, whether speech or music, strengthens learned identities in its acoustic space and enhances categorical processing (Bidelman and Lee, 2015; Iverson et al., 2003; Kuhl, 2004; Moon et al., 2013).

Human neurophysiological recordings (M/EEG, fMRI) have revealed a distributed fronto-temporo-parietal neural network supporting auditory categorization including bilateral superior temporal gyrus (STG), inferior parietal, motor cortex, and prefrontal regions (e.g., Alho et al., 2016; Bidelman and Lee, 2015; Binder et al., 2004; Chang et al., 2010; Feng et al., 2018; Golestani et al., 2002; Golestani and Zatorre, 2004; Lee et al., 2012; Liebenthal et al., 2010; Luthra et al., 2019; Myers and Blumstein, 2008; Myers et al., 2009). Yet, specific recruitment of these areas is modulated by task demands (Feng et al., 2018), item feedback (Yi and Chandrasekaran, 2016) and attention (Bidelman and Walker, 2017), whether categorization is rule-based or implicit (Yi et al., 2016), stimulus familiarity (e.g., native vs. nonnative speech: Bidelman and Lee, 2015), and presumably, the acoustic domain in which behavior is operating (e.g., speech vs. music). Particularly important to CP is a strong neural interface between temporal and frontal cortices (Bizley and Cohen, 2013; Blumstein et al., 1977; Chevillet et al., 2013; DeWitt and Rauschecker, 2012; Jiang et al., 2018; Luthra et al., 2019). Still, neuroimaging work is equivocal on the locus of categorical (perceptually invariant) brain representations with regard to auditory-linguistic hubs of the CP network. Intracranial recordings suggest abstract speech categories arise within primary auditory cortex (PAC) (Bouton et al., 2018; Chang et al., 2010) whereas fMRI implicates left inferior frontal gyrus (IFG) in categorical formation (Lee et al., 2012; Myers et al., 2009). A possible reconciliation of divergent findings may be that PAC and IFG are differentially engaged during CP depending on intrinsic and extrinsic influences (e.g., stimulus factors; lexical competition; listeners’ experience) (e.g., Bidelman and Lee, 2015; Luthra et al., 2019). Under investigation here is whether PAC-IFG pathways underlying auditory categorical decisions might be shaped and even differentially engaged in an experience-dependent manner. That is, we asked whether categories are formed within *different* auditory-linguistic brain regions in highly skilled vs. novice perceivers.

To this end, we took a neuroethological approach (Suga, 1989) to investigate the neural mechanisms underlying auditory categorization by examining individuals with highly exaggerated and specialized listening abilities: musicians. Musicians represent an ideal human model to understand experience-dependent plasticity in auditory perceptual-cognitive functions (Herholz and Zatorre, 2012; Kraus and Chandrasekaran, 2010; Moreno and Bidelman, 2014; Munte et al., 2002; Zatorre and McGill, 2005). Germane to the current study, we recently demonstrated trained musicians have enhanced (faster and more discrete) categorization of speech sounds compared to their nonmusician peers (Bidelman, 2017; Bidelman and Alain, 2015; Bidelman et al., 2014). These behavioral benefits were accompanied by electrophysiological enhancements in auditory encoding 150-200 ms after sound onset. While the underlying locus of these neuroplastic effects (observed in scalp EEG) has yet to be defined, their early latency (< 200 ms) strongly implies categorical representations might exist as early as PAC (e.g., Bidelman and Lee, 2015; Chang et al., 2010), at least in highly trained listeners (i.e., musicians).

To probe these questions, we recorded high-density neuroelectric brain activity (EEG) in musicians and nonmusicians while they rapidly categorized speech and musical sounds. Source reconstruction and functional connectivity analyses parsed the underlying brain mechanisms of categorization and differential engagement of auditory-linguistic hubs of the CP network depending on experience. Following notions that musicianship expands speech selective brain regions (Dick et al., 2011) and automatizes categorical processing (Bidelman and Alain, 2015; Bidelman et al., 2014; Elmer et al., 2012), we hypothesized early auditory cortical representations (i.e., PAC) would suffice categorization in highly trained listeners. Moreover, if PAC representations are weaker (less categorically organized) in unskilled listeners, we further predicted nonmusicians would require additional recruitment of IFG to enable successful categorization. Domain-specificity was tested by comparing neurobehavioral responses to speech vs. music. Our findings reveal a double dissociation in functional recruitment of the auditory-linguistic network (PAC-IFG) subserving CP that depends on musicianship and stimulus domain. Whereas highly experienced listeners (musicians) show categorical neural organization in early PAC, inexperienced listeners (nonmusicians) must broadcast information to higher-order linguistic brain areas (IFG) in order to generate category representations.

## MATERIALS & METHODS

### Participants

Twenty young adults participated in the experiment: 10 musicians (6 female) and 10 nonmusicians (9 female). All reported normal hearing sensitivity (i.e., ≤ 25 dB HL; 500-4000 Hz) and no history of neuro-psychiatric illness. Musicians (M) were defined as amateur instrumentalists with ≥ 8 years of continuous private instruction on their principal instrument (*mean* ± *SD*; 15.0 ± 6.2 yrs), beginning prior to age 12 (6.9 ± 3.2 yrs). Nonmusicians (NM) had <2 years of lifetime music training (0.61 ± 0.85 yrs). These inclusion criteria are consistent with previous reports on musicianship and neuroplasticity (Mankel and Bidelman, 2018; Parbery-Clark et al., 2009; Wong et al., 2007; Yoo and Bidelman, 2019; Zendel and Alain, 2009). All but one participant was right-handed (Oldfield, 1971) and all were native speakers of American English. The two groups were otherwise matched in age (M: 22.4 ± 4.5 yrs, NM: 22.5 ± 2.8 yrs; *t*_18_ = −0.06, *p* = 0.95), years of formal education (M: 16.6 ± 3.3 yrs, NM: 17.1 ± 2.2 yrs; *t*_18_ = - 0.04, *p* = 0.97), and gender balance (Fisher exact test, *p*=0.30). Participants were paid for their time and gave written informed consent in compliance with a protocol approved by the University of Memphis IRB.

### Stimuli

We used speech and music continua from our previous reports on the neural mechanisms of CP (e.g., Bidelman and Alain, 2015; Bidelman et al., 2013; Bidelman and Walker, 2017; Bidelman et al., 2014).

#### Speech continuum

Speech tokens comprised a five-step synthetic vowel continuum (Bidelman et al., 2013) (**Fig. 1a**). Tokens were 100 ms (10 ms ramps). Each contained an identical voice fundamental (F0), second (F2), and third formant (F3) frequencies (F0: 150, F2: 1090, and F3: 2350 Hz). F1 was parameterized over five equidistant steps (430 to 730 Hz) resulting in perceptual phonetic continuum from /u/ to /a/.

**Figure 1:**
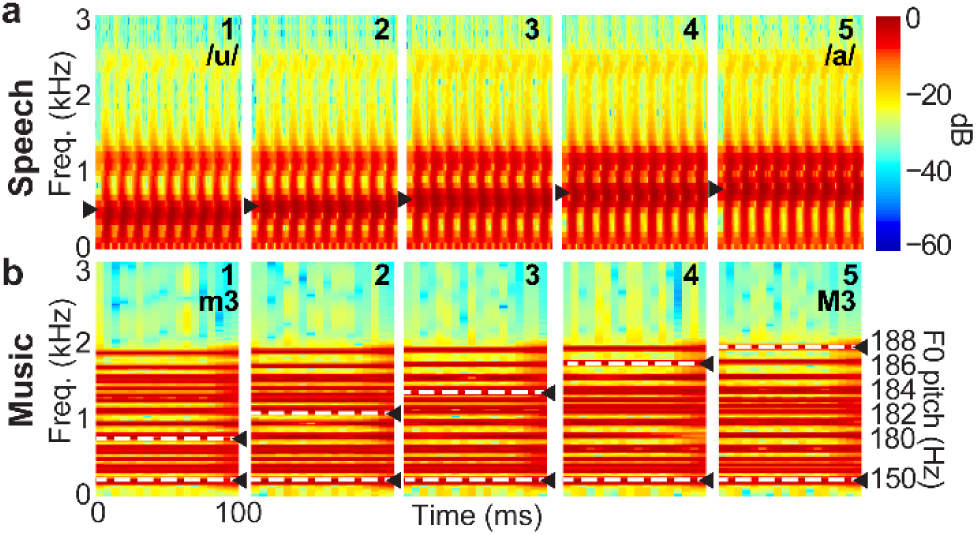
Spectrograms of speech and music continua used to probe experience-dependent plasticity in CP. (**a**) For speech, vowel first formant frequency was varied across five equal steps (430-730 Hz; ▸) creating a continuum from /u/ to /a/. (**b**) For music, tone complexes formed two-tone pitch intervals (white dotted lines) spanning a continuum from a minor (m3) to major (M3) third. Stimuli were otherwise matched in duration (100 ms), intensity (83 dB SPL), and starting pitch height (F0=150 Hz).

#### Music continuum

We used a comparable five-step continuum of pitch intervals to assess CP for music (Bidelman and Walker, 2017) (**Fig. 1b**). Individual notes were synthesized using complex-tones (10 iso-amplitude harmonics; 100 ms duration). For each token, the lower tone of the dyad was fixed with a F0 of 150 Hz (matching the F0 of the speech continuum) while the upper tone’s F0 varied over five equal steps to produce a perceptual continuum of musical intervals between the minor (m3; f_lower_ = 150, f_higher_ = 180 Hz) and major (M3; *f*_*lower*_ = 150, *f*_*higher*_ = 188 Hz) third on the chromatic scale (e.g., Burns and Ward, 1978). The m3 and M3 intervals connote the valence of “sadness” (m3) and “happiness” (M3) even to non-musicians (Brattico et al., 2009) and are easily described to listeners unfamiliar with music-theoretic labels (Bidelman and Walker, 2017).

### Task procedure

During EEG recording, listeners heard 200 randomly ordered exemplars of each speech/music token (presented in separate blocks). They were asked to label each sound with a binary response as quickly and accurately as possible (*speech*: “u” or “a”; *music*: “minor 3^rd^ or “major third”). Stimuli were delivered binaurally at 83 dB SPL through insert earphones (ER-2; Etymotic Research). The interstimulus interval (ISI) was jittered randomly between 400 and 600 ms (20 ms steps, rectangular distribution) following listeners’ behavioral response to avoid anticipating the next trial and rhythmic entrainment of the EEG.

### EEG recording and preprocessing

Continuous EEGs were recorded from 64 sintered Ag/AgCl electrodes at standard 10-10 locations around the scalp (Oostenveld and Praamstra, 2001). Recordings were sampled at 500 Hz (SynAmps RT amplifiers; Compumedics Neuroscan) and passband filtered online (DC-200 Hz). Electrodes placed on the outer canthi of the eyes and the superior and inferior orbit were used to monitor ocular movements. During acquisition, electrodes were referenced to an additional sensor placed ∼ 1 cm posterior to the Cz channel. Data were re-referenced off-line to the common average for analysis (Bertrand et al., 1985). Contact impedances were maintained < 10 kΩ during data collection.

Subsequent preprocessing was performed in Curry 7 (Compumedics Neuroscan) and BESA® Research (v7) (BESA, GmbH). Ocular artifacts (i.e., blinks and saccades) were first corrected in the continuous EEG using a principal component analysis (PCA) (Picton et al., 2000). Cleaned EEGs were then digitally filtered (1-30Hz; zero-phase filters), epoched (−200-800 ms), baseline corrected to the prestimulus interval, and ensemble averaged to derive responses for each stimulus condition per participant. This resulted in 10 ERP waveforms per participant (5 tokens × 2 stimulus domains).

### Behavioral data analysis

For each continuum, identification scores were fit with a two-parameter sigmoid function: *P* = 1/[1+*e*^-*β1*(*x* - *β0*)^], where *P* is the proportion of trials identified as a given vowel, *x* is the step number along the stimulus continuum, and *β*_*0*_ and *β*_*1*_ the location and slope of the logistic fit estimated using nonlinear least-squares regression (Bidelman and Walker, 2017; Bidelman et al., 2014). These parameters were used to assess differences in the location and “steepness” (i.e., rate of change) of the categorical boundary as a function of stimulus domain (i.e., speech vs. music) and group (M vs. NM). Larger *β*_*1*_ values reflect steeper psychometric functions and hence, indicate stronger CP. Behavioral speech labeling speeds (i.e., reaction times; RTs) were computed as listeners’ median response latency across trials for a given condition. RTs outside 250-2500 ms were deemed outliers and excluded from further analysis (Bidelman et al., 2013; Bidelman and Walker, 2017).

### Electrophysiological data analysis

#### Source reconstruction

Sensor (electrode)-level recordings were transformed to source space using discrete inverse models to directly assess the neural generators underlying experience-dependent plasticity in CP. We used Classical Low Resolution Electromagnetic Tomography Analysis Recursively Applied (CLARA) [BESA (v7)] (Iordanov et al., 2014) to estimate the neuronal current density underlying the scalp potentials for speech (e.g., Alain et al., 2017; Bidelman et al., 2018). CLARA models the inverse solution as a large collection of elementary dipoles distributed over nodes on a mesh of the cortical volume. The algorithm estimates the total variance of the scalp-recorded data and applies a smoothness constraint to ensure current changes minimally between adjacent brain regions (Michel et al., 2004; Picton et al., 1999). CLARA renders more focal source images by iteratively reducing the source space during repeated estimations. On each iteration (x3), a spatially smoothed LORETA solution (Pascual-Marqui et al., 2002) was recomputed and voxels below a 1% max amplitude threshold were removed. This provided a spatial weighting term for each voxel on the subsequent step. Three iterations were used with a voxel size of 7 mm in Talairach space and regularization (parameter accounting for noise) set at 0.01% singular value decomposition. Group-level statistical (*t*-stat) maps were computed using the ‘ft_sourcestatistics’ function in the MATLAB FieldTrip toolbox (Oostenveld et al., 2011) and threshold at α=0.05. Source activations were then visualized by projecting them onto the semi-inflated MNI adult brain template (Fonov et al., 2009).

From each CLARA volume (i.e., activation timecourse per voxel), we extracted the amplitude of source activity in predefined regions of interest (ROIs) including bilateral primary auditory cortex (PAC) and inferior frontal gyrus (IFG) near Broca’s area (see Fig. 4). These ROIs were selected given their known role in complex speech perception including auditory categorization (e.g., Alain et al., 2018; Bidelman et al., 2018; Bidelman and Howell, 2016; Bizley and Cohen, 2013; Du et al., 2014; Mazziotta et al., 1995; Scott and Johnsrude, 2003). The spatial resolution of CLARA is 5-10 mm (Iordanov et al., 2016; Iordanov et al., 2014), which is considerably smaller than the distance to resolve PAC and IFG (∼40 mm; Mazziotta et al., 1995).

To quantify the degree to which neural responses showed categorical coding, we averaged source amplitudes to prototypical tokens at the ends of the continua and compared this combination to the ambiguous token at its midpoint (e.g., Bidelman, 2015; Bidelman and Walker, 2017; Liebenthal et al., 2010). This contrast (i.e., mean[Tk1, Tk5] vs. Tk 3) allowed us to assess the degree to which each groups’ neural activity differentiated stimuli with well-formed categories from those heard with a bistable (ambiguous) identity within the speech and music domains.

To quantify brain-behavior relationships, we extracted the overall response amplitude within the PAC and IFG ROI centroids, computed at the latency of maximum global field power (GFP) on the scalp (Lehmann and Skrandies, 1980) (see Fig. 4a). Peak GFP occurred at 288 ms for speech and 226 ms for music. We used Spearman correlations to assess if PAC vs. IFG amplitudes predicted the slopes of listeners’ psychometric functions when categorizing stimuli per domain (speech, music). This analysis was repeated for both left and right hemispheres to test whether lateralized activity was more predictive of behavior in a certain domain (e.g., right PAC driving music CP; left PAC driving speech CP).

#### Functional connectivity

We measured causal (directed) flow of information within the auditory-linguistic brain network using Granger Causality (GC) (Geweke, 1982; Granger, 1969). Functional connectivity was computed in BESA Connectivity (v1), which computes GC in the frequency domain (Geweke, 1982) using non-parametric spectral factorization on single-trial time-frequency maps (Dhamala et al., 2008). The frequency decomposition was based on complex demodulation (Papp and Ktonas, 1977), akin to a short-term (running) Fourier transform, that provides uniform frequency resolution across the bandwidth of analysis. Signal *X* is said to “Granger-cause” signal *Y* if past values of *X* contain information that predict *Y* above and beyond information contained in past values *Y* alone. Importantly, GC can be computed directionally (e.g., *X→Y*) to infer causal flow between interacting brain signals. We computed GC between PAC→IFG activity using full-band (1-30 Hz) responses at the latency corresponding to the max GFP (see Fig. 4a), where neural responses showed group differences in the CLARA maps. Connectivity was only computed for the Tk1/5 tokens to avoid neural signaling that might be related to resolving ambiguous (bistable) sounds (e.g., Tk 3) and because these tokens elicited stronger responses than the midpoint tokens (i.e., Tk1/5 > Tk3; Fig. 4b).

## RESULTS

### Behavioral categorization (% and RTs)

**Figure 2** shows behavioral psychometric identification functions for Ms and NMs when classifying speech (**Fig. 2a**) and music (**Fig. 2b**). Listeners’ identification was generally dichotomous as indicated by an abrupt shift in perception midway through the continua. For NMs, music stimuli elicited more continuous perception as indicated by the lack of any abrupt perceptional shift and linear/flat psychometric function. Both Ms and NMs showed stronger CP for speech than music [*M*: *t*_18_ = 2.38, *p* = 0.028; *NM*: *t*_18_ = 6.36, *p*<0.0001]. However, Ms demonstrated considerably sharper perceptual boundaries than NMs for both speech [*β*_*1*_ parameter; *t*_18_ = 2.23, *p* = 0.034] and music [*t*_18_ = 2.85, *p* = 0.011] continua. These findings suggest that while CP is stronger for speech than musical sounds, musicians show enhanced perceptual categorization in both auditory domains.

Behavioral RTs for speech and music categorization are shown for each group in **Figure 2c-d.** An ANOVA conducted on speech labeling speeds revealed RTs were modulated across vowel token [*F*_4, 72_ = 32.60, *p*<0.0001]. There was no effect of group [*F*_1, 18_ = 1.17, *p*=0.29] which might be expected given the overlearned nature of vowels in our native English speakers. Still, both Ms [*t*_72_= 9.12, *p*<0.0001] and NMs [*t*_*72*_= 6.11, *p*<0.0001] showed the characteristic slowing near the CP boundary (token 3) relative to other tokens along the continuum [i.e., mean(Tk1,2,4,5) < Tk3], consistent with previous reports examining speeded speech classification (Bidelman et al., 2013; Bidelman and Walker, 2017; Pisoni and Tash, 1974). For music, we found a group x token interaction [*F*_4, 72_ = 2.59, *p*=0.043]. Interestingly, Ms showed a categorical slowing in RTs near the ambiguous midpoint [*t*_*72*_= 2.05, *p*=0.044] but this bowing effect was not observed in NMs [*t*_*72*_= −1.33, *p*=0.19]. This again suggests music was perceived less categorically in NMs compared to trained Ms.

**Figure 2:**
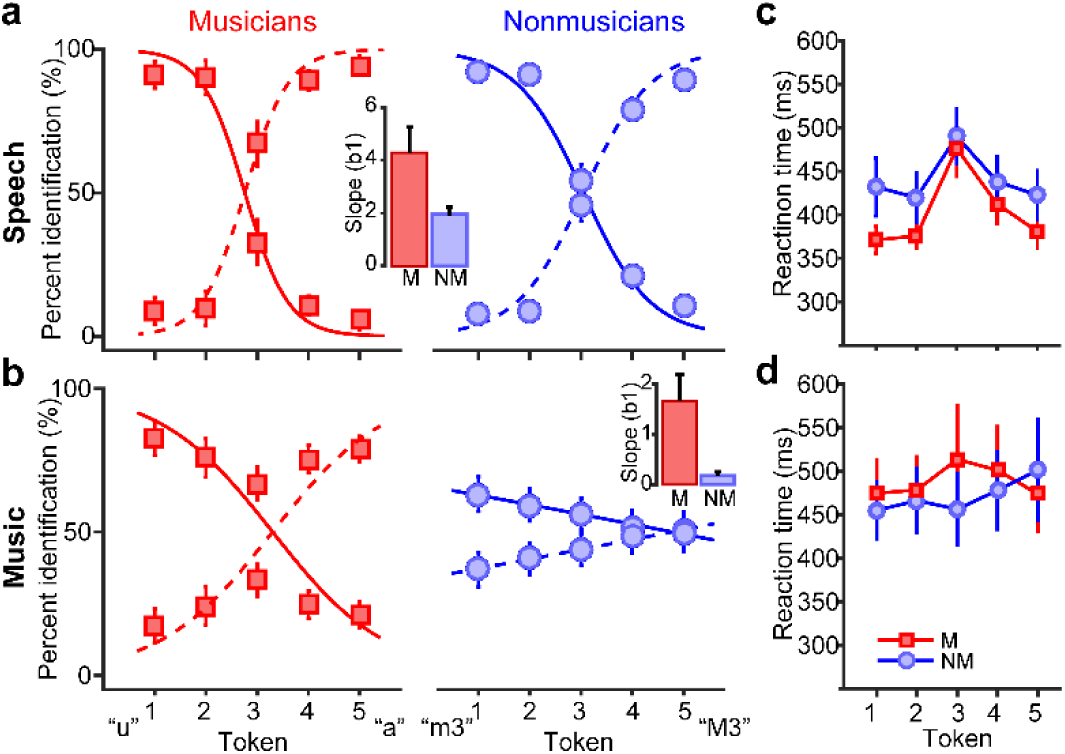
Perceptual categorization for speech and music is enhanced in musicians. (**a**) Psychometric identification functions for speech show an abrupt shift in behavior indicative of discrete CP. (**b**) Music identification is discrete (categorical) in Ms but continuous in NMs. Ms show stronger CP than NMs (steeper identification curves) in both domains (*insets*). (**c-d**) Reaction times for classifying stimuli. Listeners are slower to label speech near the categorical boundary (Tk 3), indicative of CP (Bidelman et al., 2013; Pisoni and Tash, 1974) but Ms are faster at making categorical judgments across the board. Ms show a similar categorical speed effect (bowing in RTs) for music that is not observed in NMs. errorbars = ± 1 s.e.m.; M, musicians; NM, nonmusicians.

### Electrophysiological data

Butterfly ERP plots (sensor-level potentials) are shown per group and stimulus domain in **Figure 3**. Visual inspection revealed more robust cortical responses in Ms compared to NMs for both speech and musical sounds, particularly at fronto-central electrodes where CP effects appear most prominent on the scalp surface (e.g., Bidelman and Walker, 2017; Liebenthal et al., 2010). The latency of these effects (∼250 ms) corresponds roughly to the P2 wave, a deflection previously suggested as an electrophysiological marker of CP (Bidelman and Alain, 2015; Bidelman et al., 2013; Liebenthal et al., 2010).

**Figure 3:**
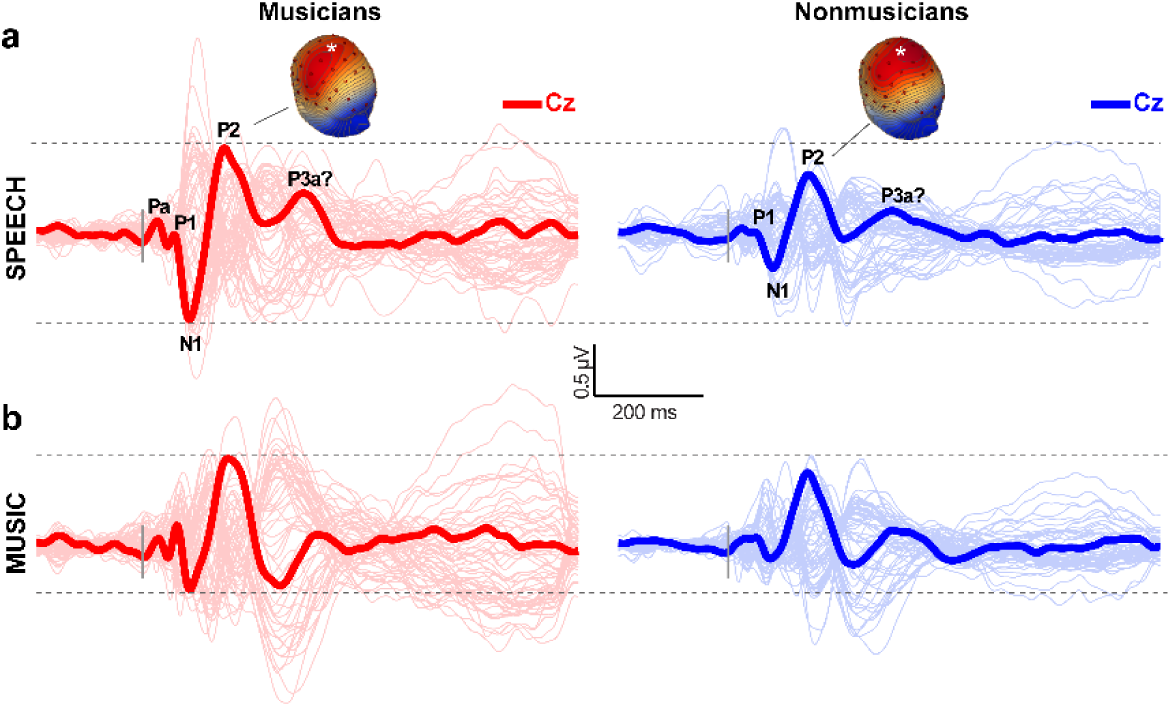
Scalp-recorded event-related brain potentials (ERPs) show stronger neural responses to speech and music in musicians. Grand averaged butterfly plot of neuroelectric time waveforms per group and stimulus domain. Cortical ERPs appear as biphasic deflections (e.g., P1-N1-P2 “waves”) within ∼200 ms after the time-locking stimulus. Cz electrode = bold trace. Neural activity is modulated by group and stimulus domain. Scalp topographies are plotted at the latency of the P2 wave (*Cz electrode). Vertical bars=stimulus onset (*t*=0). Dotted lines, visual aid for group comparisons.

**Figure 4:**
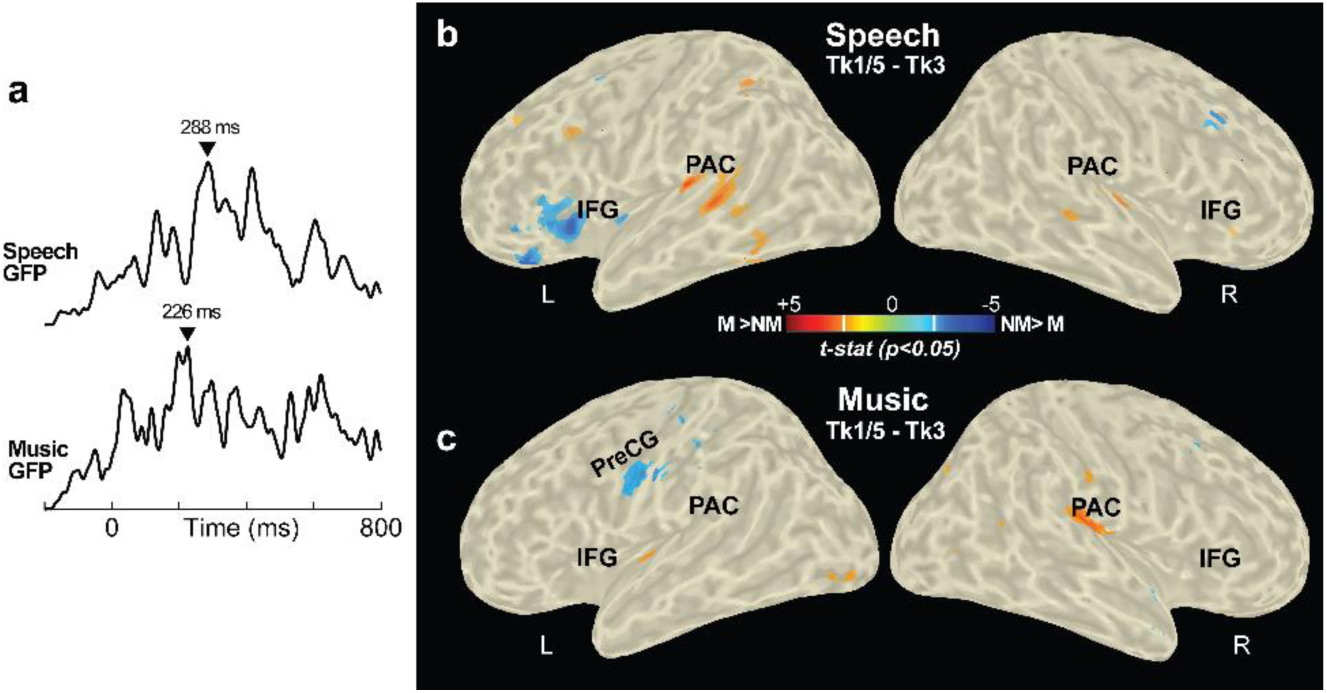
Source responses reveal differential activation of auditory and linguistic cortex during categorization depending on musical training. (**a**) Grand average global field power (Lehmann and Skrandies, 1980) (collapsed across tokens and groups) shows the time-course of aggregate neural activity across the scalp for each domain. (**b-c**) Statistical contrast maps of CLARA activations between musicians and nonmusicians. Maps are shown at the maximum GFP latency (▾, Fig. 4a), projected onto the semi-inflated MNI adult brain template (Fonov et al., 2009) and contrast group differences (*t*-stat, *p* < 0.05 masked, uncorrected) in the degree of categorical coding (i.e., Tk1/5 – Tk3 effect; Bidelman and Walker, 2017) across the entire brain volume. (**b**) For speech, categorical coding is observed in bilateral PAC for Ms but left IFG for NMs. (**c**) For music, Ms show stronger categorical responses than NMs in right PAC; NMs show stronger categorization in left preCG (motor cortex). PAC, primary auditory cortex; IFG; inferior frontal gyrus; preCG, precentral gyrus (primary motor cortex); L/R, left/right hemisphere.

CLARA imaging parsed the underlying neural mechanisms responsible for M’s enhanced CP and these electrode-level effects (**Fig. 4**). Across all tokens and groups, neural activity peaked ∼50 ms earlier than speech (GFP_speech_ latency: 288 ms; GFP_music_ latency: 226 ms; **Fig. 4a**). To quantify the degree of categorical neural coding, we averaged source amplitudes to prototypical tokens at the end of the continua and compared this combination to the ambiguous midpoint token (e.g., Bidelman, 2015; Bidelman and Walker, 2017; Liebenthal et al., 2010). This contrast (i.e., mean[Tk1, Tk5] vs. Tk 3) allowed us to assess when and where neural responses (CLARA maps) differentiated stimuli with well-formed categories from those heard with a bistable (ambiguous) identity. Contrasts of this categorical coding effect revealed a double-dissociation within auditory-linguistic brain regions depending on group and stimulus domain (**Figs. 4 b,c**). For speech, musicians showed stronger categorical responses in PAC bilaterally, whereas in nonmusicians, categorical speech responses were observed primarily in left IFG. For music, Ms showed stronger categorical responses than NMs in right PAC. In contrast, categorical coding for music was stronger in NMs in left precentral gyrus (motor cortex), a region previously identified to predict CP for speech (Chevillet et al., 2013). These results suggest a differential engagement of the auditory-linguistic-motor loop during auditory categorization depending on musicianship.

Brain-behavior correlations again revealed a double-dissociation in the predictive power of PAC vs. IFG in predicting behavior. Across cohorts, larger categorical speech activity (Tk1/5 > Tk3) in left PAC (as in Ms) was associated with more dichotomous CP (steeper psychometric slopes) [*r*=0.46, *p*=0.042] (**Fig. 5b**). We found the reverse in left IFG which negatively correlated with behavior [*r*=-0.68, *p*=0.0009] (**Fig. 5a**), where stronger neural activity predicted poorer perception. Neither right hemisphere ROI nor any of the music responses correlated with behavior (all *p*s > 0.27; data not shown).

**Figure 5:**
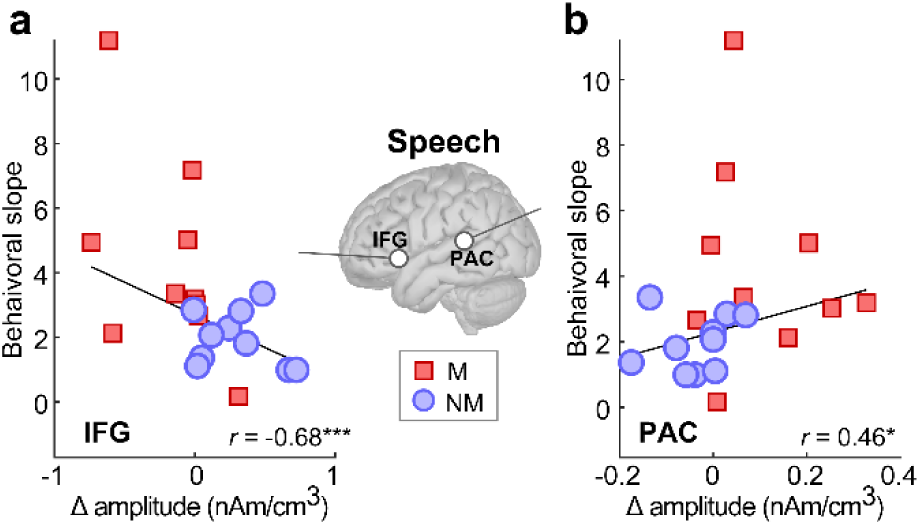
Brain-behavior correlations reveal a double-dissociation in how PAC and IFG predict behavioral CP in expert and non-expert listeners. Neural responses reflect activations to speech within PAC and IFG at the centroid of each ROI shown (see Fig. 4). MNI coordinates (*x,y,z*): PAC_LH_ = [-38.5, −37.5, 5.5], PAC_RH_= [41.5, −26.5, 0.5], IFG_LH_ =[−22.5, 24.5, −2.5], and IFG_RH_ =[23.5, 26.5, −2.5] mm. Larger differentiation of speech (i.e. Δ amplitude: Tk1/5 > Tk3) in left PAC is associated with stronger categorical percepts (steeper psychometric slopes). The reverse is observed in left IFG, where stronger activity predicts poorer behavior. Neither right hemisphere PAC/IFG nor music responses correlated with behavior (data not shown). \**p*<0.05, \*\*\**p*<0.001.

Increased involvement of IFG in nonmusicians could reflect the need to recruit additional higher-order (linguistic) brain areas downstream if neural representations in auditory cortex are insufficient for categorization. We tested this possibility by measuring directed functional connectivity between PAC and IFG using Granger causality (GC), an information-theoretic measure of causal signal interactions (Geweke, 1982; Granger, 1969). GC values were cube-root transformed to improve homogeneity of variance assumptions for parametric statistics and to account for the lower bound of GC values (=0). A three-way, mixed-model ANOVA (group x stimulus domain x hemisphere; subject=random) revealed a significant group x stimulus interaction [*F*_1, 36_ = 6.89, *p*=0.0126], meaning the strength of PAC-IFG connectivity was modulated dependent on listeners’ musical training and whether they were categorizing speech or music. By hemisphere, we found a stimulus x group effect in LH [*F*_1, 36_ = 4.76, *p*=0.0358] (**Fig. 6a**). Tukey-Kramer comparisons revealed the LH interaction was attributable to stronger connectivity for speech than music in NMs, whereas musicians’ LH PAC-IFG connectivity was equally strong between stimulus domains (i.e., *NMs*: speech > music; *Ms*: speech=music). By domain, contrasts showed marginally stronger PAC-IFG connectivity in NMs vs. Ms for speech (*p*=0.08) but not music (*p*=0.26). In contrast, no effects in RH were significant (all *p*s > 0.122) (**Fig. 6b**). These findings confirm a differential engagement of the left-lateralized auditory-linguistic network (PAC-IFG) between musicians and nonmusicians during auditory categorization that also depends on stimulus domain.

**Figure 6:**
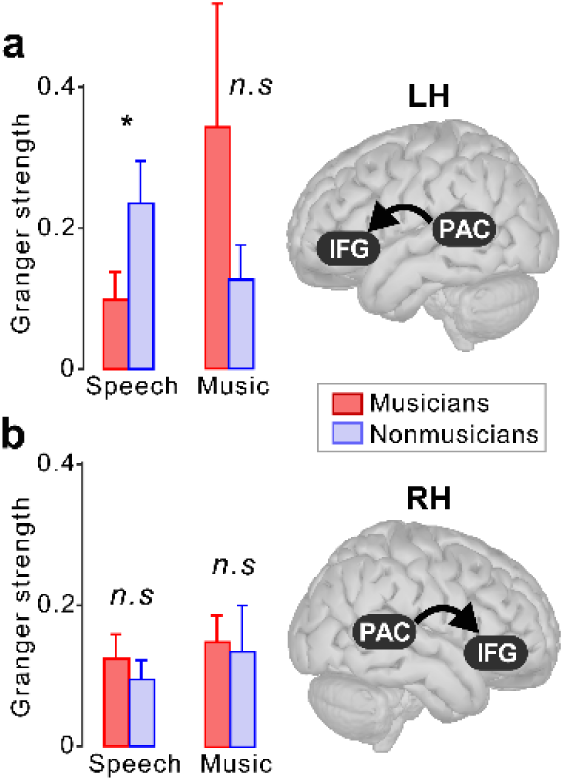
Functional connectivity reveals experience-dependent changes in causal neural signaling within the auditory-linguistic pathway. **(a)** Granger causality between PAC directed toward IFG shows stronger communication during speech compared to music categorization in NMs. Contrastively, left PAC-IFG connectivity is invariant across stimulus domains in Ms. (**b**) PAC-IFG connectivity does not vary in RH. These results suggest a differential engagement of the left-lateralized auditory-linguistic network (PAC-IFG) depending on music expertise. Whereas neural representations in PAC are sufficient for categorization in Ms (see Fig. 4), NMs recruit additional linguistic areas downstream to aid perception. \**p*<0.05.

## DISCUSSION

By measuring electrical brain activity in highly specialized (musicians) vs. novice (nonmusicians) listeners we demonstrate that categorical processing for speech and musical sounds varies in an experience-dependent manner. Our EEG data reveal the neural mechanisms underlying these behavioral enhancements are accompanied by different engagement and neural signaling within the auditory-linguistic pathway (PAC-IFG) that critically depend on listening experience. Whereas highly skilled musicians code auditory categories in primary auditory cortex (PAC), nonmusicians show additional recruitment of non-auditory regions (e.g., inferior frontal gyrus, IFG) to successfully generate category-level information. Our findings provide new evidence that the brain arrives at categorical labels through different operational “modes” within an identical PAC-IFG pathway. Whereas early canonical PAC representations are sufficient to generate categories in highly trained ears, less experienced listeners must broadcast information downstream to non-canonical, higher-order linguistic brain areas (IFG) to construct abstract sound labels.

Previous work has shown musicianship enhances the ability to categorize musically-relevant sounds, e.g., pitch intervals and chords (Burns and Ward, 1978; Howard et al., 1992; Klein and Zatorre, 2011; Locke and Kellar, 1973; Siegel and Siegel, 1977; Zatorre and Halpern, 1979). Our data confirm enhanced music categorization in musicians but further extend prior studies by demonstrating that training also enhances the perceptual categorization of speech (cf. Bidelman and Alain, 2015; Bidelman et al., 2014; Elmer et al., 2012). Musicians were faster at categorizing speech and music and achieved more pronounced (i.e., steeper) identification, indicating musicianship enhances sound-to-meaning relations in a domain general manner. These findings support notions that music expertise warps or restricts the perceptual space near category boundaries, sharpening the internal representations of auditory categories and supplying more behaviorally-relent decisions when classifying sound objects (Bidelman and Alain, 2015; Bidelman et al., 2014). Moreover, comparisons across stimulus domains indicated less categorical (more continuous) music perception in NM listeners. Nonmusicians’ weaker CP for music can be parsimoniously explained as an unfamiliarity in associating verbal labels to the pitch intervals of music. Just as language experience sharpens categories for a speaker’s native speech sounds (Bidelman and Lee, 2015; Bradlow et al., 1997; Kuhl et al., 1992; Xu et al., 2006), we find musicians (but not NMs) perceive music categorically. Our findings bolster notions that less familiar sounds not encountered in one’s regular experience fail to perceptually organize in a categorical manner (e.g., Bidelman and Lee, 2015; Bidelman and Walker, 2017; Burns and Campbell, 1994; Burns and Ward, 1978; Xu et al., 2006; Zatorre, 1983). More broadly, these plasticity data counter universalist views of CP (Harnad, 1987) that suggest categorical boundaries are somehow innate (e.g., innate-sensitivity hypothesis; Rosen and Howell, 1987) or occur naturally due to acoustic and/or neural discontinuities imposed by auditory system constraints (Berlin and Kay, 1969; Harnad, 1987; Miller et al., 1976) by instead revealing a strong role of experience in shaping categorical organization.

Our EEG data corroborate behavioral findings by unmasking the underlying brain mechanisms driving experience-dependent changes in auditory categorization. We found musicians had stronger neural encoding of speech and music stimuli across the board (Fig. 3), consistent with previous neuroimaging studies (Bidelman and Alain, 2015; Bidelman et al., 2014; Musacchia et al., 2008; Schneider et al., 2002; Shahin et al., 2003; Wong et al., 2007). These results support growing evidence that musicians’ brain responses carry more information relevant to perception than their non-musician counterparts (Bidelman et al., 2014; Elmer et al., 2012; Weiss and Bidelman, 2015). However, we found stark differences in the underlying brain regions mediating each group’s CP. Whereas skilled listeners (musicians) were able to code auditory categories in PAC, naïve ears (nonmusicians) showed categorical responses in non-canonical areas including IFG. This double-dissociation in source contribution between groups suggests that listening experience differentially shapes the functional coupling within the auditory-linguistic network.

There is evidence that sound representations in PAC might self-organize due to non-uniformities in cell firing between exemplar vs. non-exemplar sounds (cf. within vs. between category tokens) (Guenther and Gjaja, 1996). Receptive fields of auditory cortical neurons also show marked changes in their temporal discharge patterns across categorically perceived speech continua (Steinschneider et al., 2003; Steinschneider et al., 1999). Therefore, one interpretation of our data is that musical training enhances CP by automatizing the categorization process (i.e., binning sounds into perceptually-relevant groupings) and relegating category-level representations to the earliest stages of auditory cortical processing. The fact that we find strong categorical organization in PAC only among musicians supports this notion. On the contrary, PAC representations are insufficient for categorical organization in nonmusicians, and these less skilled listeners must compute categories in higher-order frontal regions (IFG) downstream. Higher fidelity auditory neural representations local to PAC would tend to speed behavioral decisions and result in steeper, more discrete identification (Bidelman et al., 2019; Bidelman et al., under review; Rozsypal et al., 1985) as we find in our musician cohort. Alternatively, focal readout of sensory encoding (PAC) by prefrontal regions (IFG) (Binder et al., 2004; Bouton et al., 2018) may underlie the increased decision variability we find in nonmusicians.

Our data align with general notions that non-canonical auditory regions including IFG (Broca’s area) and adjacent prefrontal areas play a major role in the categorical processing (Bouton et al., 2018; Lee et al., 2012; Luthra et al., 2019; Meyers et al., 2008; Myers et al., 2009). Consistent with our findings, phoneme category selectivity is observed early (<150 ms) in both PAC and left inferior frontal gyrus (pars opercularis) (Alho et al., 2016; Bidelman et al., 2013; Chang et al., 2010; Toscano et al., 2018). Still, cue- and category-based representations are probably not mutually exclusive and may overlap in both time and space within the brain; IFG, for instance, may code acoustic details of speech nearly simultaneously (within ∼50 ms) of coding phonological categories (Toscano et al., 2018). In nonhuman primates, inferior frontal regions show categorical coding even during visual classification, suggesting a broad role of this area in creating perceptual invariance (Freedman et al., 2001). Still, our findings suggest that IFG may not play a *domain-general* role in computing category representations (Myers et al., 2009). Indeed, we found IFG activity anti-correlated with behavior for speech (Fig. 5); increased IFG activity, as in nonmusicians, was indicative of poor behavioral categorization performance. In contrast, stronger PAC responses predicted better behavior. Similar phoneme-category selectivity has also been observed in left PreCG and connectivity between posterior aspects of auditory cortex and PreCG have been shown to mediate complex speech identification decisions (Chevillet et al., 2013). In the present study, we found that nonmusicians did show categorical responsivity in PreCG, but only for music. PreCG activation in nonmusicians could reflect the increased difficulty of classifying unfamiliar music tokens, which would be expected to evoke non-auditory regions to aid perception (Du et al., 2014). However, our data do not readily support interpretations that sensorimotor interplay within the dorsal auditory-motor stream is sufficient to account for performance in categorization tasks (cf. Chevillet et al., 2013). Instead, we find engagement of non-canonical areas (e.g., PreCG, IFG) outside of auditory system depends critically on both what an observer is classifying (e.g., speech vs. music) and their relative experience with the sounds of that domain.

While IFG has been implicated in speech identification and categorization (Du et al., 2014; Lee et al., 2012; Luthra et al., 2019), nonmusicians’ increased frontal activation could instead be related to increased attentional load (Bouton et al., 2018; Giraud et al., 2004), lexical uncertainty (Luthra et al., 2019), and/or unconscious sensory repair that applies prior knowledge to a noisy input (Shahin et al., 2009). Indeed, decision loads IFG during effortful speech listening (Binder et al., 2004; Bouton et al., 2018; Du et al., 2014) and left IFG is more active when items are perceptually confusable (Feng et al., 2018). Therefore, insomuch as nonmusicians are less skilled listeners and were more challenged by even our simple auditory CP tasks (as suggested by their increased RTs), IFG might necessarily be recruited in a top-down manner to aid categorical predictions on the sensory input. Supporting this interpretation, studies on categorical speech learning have shown frontal speech regions are relatively less active in “good learners” (Golestani and Zatorre, 2004), mirroring our findings in musicians. Alternatively, IFG regions may reflect subvocal rehearsal strategies or articulatory codes that aid speech perception (Du et al., 2014; Golestani and Zatorre, 2004; Hickok and Poeppel, 2004; Papoutsi et al., 2009). Functional coupling between auditory and frontal brain regions is modulated by musical training (current study; Du and Zatorre, 2017). Thus, it stands to reason that nonmusicians’ increased involvement of IFG may reflect augmented generation or retrieval of articulatory codes to improve speech categorization. Similarly, the lack of IFG involvement for music may reflect total absence of verbal labels of articulatory code for non-speech stimuli and/or NM’s substantial difficulty in the music task (Fig. 2).

Our data further align with predictive coding models of speech processing (Bouton et al., 2018; Di Liberto et al., 2018). Under this premise, more ambiguous acoustic information might invoke the need to pass information to higher-order frontal areas (IFG) to help interpret weaker sensory representations in PAC in a predictive framework. Our functional connectivity analysis supports this notion (at least for speech) as evidenced by nonmusicians’ stronger PAC→IFG neural signaling in left hemisphere. These results indicate that auditory expertise shapes not only regional engagement but also functional interplay (directed communication) between PAC and IFG when making sound-to-label associations (Bizley and Cohen, 2013; Luthra et al., 2019). They also corroborate previous studies suggesting music training enhances synchronization between auditory cortices (Kuhnis et al., 2014) and increases global network efficiency between auditory and frontal regions (Du and Zatorre, 2017; Paraskevopoulos et al., 2017).

It is worth noting that some developmental learning disorders including dyslexia have been linked to poorer categorization skills (Calcus et al., 2016; Hakvoort et al., 2016; Messaoud-Galusi et al., 2011; Noordenbos and Serniclaes, 2015; Zoubrinetzky et al., 2016). Yet, recent longitudinal (6 mo) training studies have shown improvements in categorical processing in dyslexic children in the form of stronger pre-attentive differentiation of speech at post-test (Frey et al., 2019). Such longitudinal interventions are consistent with our cross-sectional observations (present study; Bidelman and Alain, 2015; Bidelman et al., 2014), which imply active music training might fortify complex auditory processing and in turn, enhance the encoding and decoding of sound-to-meaning relations. Our study provides direct neurobiological evidence that musicianship impacts the neural architecture supporting critical and fundamental skills that support the auditory perceptual organization and categorization of sound. Consequently, it is conceivable that music rehabilitation programs might offer a means to preserve and/or offset processing difficulties in children with dyslexia (Frey et al., 2019) or those stemming from aberrant cognitive aging (Bidelman and Alain, 2015).

## Acknowledgements

Requests for data and materials should be directed to G.M.B. This work was supported by the GRAMMY Foundation^®^ and the National Institutes of Health (NIH/NIDCD R01DC016267) awarded to G.M.B.

## References

Alain, C., Arsenault, J.S., Garami, L., Bidelman, G.M., Snyder, J.S., 2017. Neural correlates of speech segregation based on formant frequencies of adjacent vowels. Scientific Reports 7, 1–11.

Alain, C., Du, Y., Bernstein, L.J., Barten, T., Banai, K., 2018. Listening under difficult conditions: An activation likelihood estimation meta-analysis. Human Brain Mapping 39, 2695–2709.

Alho, J., Green, B.M., May, P.J.C., Sams, M., Tiitinen, H., Rauschecker, J.P., Jääskeläinen, I.P., 2016. Early-latency categorical speech sound representations in the left inferior frontal gyrus. Neuroimage 129, 214–223.

Berlin, B., Kay, P., 1969. Basic Color Terms, Berkeley, CA.

Bertrand, O., Perrin, F., Pernier, J., 1985. A theoretical justification of the average reference in topographic evoked potential studies. Electroencephalography and Clinical Neurophysiology 62, 462–464.

Bidelman, G.M., 2015. Induced neural beta oscillations predict categorical speech perception abilities. Brain and Language 141, 62–69.

Bidelman, G.M., 2017. Amplified induced neural oscillatory activity predicts musicians’ benefits in categorical speech perception. Neuroscience 348, 107–113.

Bidelman, G.M., Alain, C., 2015. Musical training orchestrates coordinated neuroplasticity in auditory brainstem and cortex to counteract age-related declines in categorical vowel perception. Journal of Neuroscience 35, 1240 –1249.

Bidelman, G.M., Bush, L.C., Boudreaux, A.M., 2019. The categorical neural organization of speech aids its perception in noise. bioRxiv [preprint], doi: https://doi.org/10.1101/652842.

Bidelman, G.M., Davis, M.K., Pridgen, M.H., 2018. Brainstem-cortical functional connectivity for speech is differentially challenged by noise and reverberation. Hearing Research 367, 149–160.

Bidelman, G.M., Howell, M., 2016. Functional changes in inter-and intra-hemispheric auditory cortical processing underlying degraded speech perception. Neuroimage 124, 581–590.

Bidelman, G.M., Lee, C.-C., 2015. Effects of language experience and stimulus context on the neural organization and categorical perception of speech. Neuroimage 120, 191–200.

Bidelman, G.M., Moreno, S., Alain, C., 2013. Tracing the emergence of categorical speech perception in the human auditory system. Neuroimage 79, 201–212.

Bidelman, G.M., Sigley, L., Lewis, G., under review. Acoustic noise and vision differentially warp speech categorization.

Bidelman, G.M., Walker, B., 2017. Attentional modulation and domain specificity underlying the neural organization of auditory categorical perception. European Journal of Neuroscience 45, 690–699.

Bidelman, G.M., Weiss, M.W., Moreno, S., Alain, C., 2014. Coordinated plasticity in brainstem and auditory cortex contributes to enhanced categorical speech perception in musicians. European Journal of Neuroscience 40, 2662–2673.

Binder, J.R., Liebenthal, E., Possing, E.T., Medler, D.A., Ward, B.D., 2004. Neural correlates of sensory and decision processes in auditory object identification. Nature Neuroscience 7, 295–301.

Bizley, J.K., Cohen, Y.E., 2013. The what, where and how of auditory-object perception. Nature Reviews Neuroscience 14, 693–707.

Blumstein, S.E., Baker, E., Goodglass, H., 1977. Phonological factors in auditory comprehension in aphasia. Neuropsychologia 15, 19–30.

Bouton, S., Chambon, V., Tyrand, R., Guggisberg, A.G., Seeck, M., Karkar, S., van de Ville, D., Giraud, A.L., 2018. Focal versus distributed temporal cortex activity for speech sound category assignment. Proceedings of the National Academy of Sciences of the United States of America 115, E1299–e1308.

Bradlow, A.R., Pisoni, D.B., Yamada, R.A., Tohkura, Y., 1997. Training the Japanese listener to identify English /r/ and /l/: IV. Some effects of perceptual learning on speech production. Journal of the Acoustical Society of America 101, 2299–2310.

Brattico, E., Pallesen, K.J., Varyagina, O., Bailey, C., Anourova, I., Jarvenpaa, M., Eerola, T., Tervaniemi, M., 2009. Neural discrimination of nonprototypical chords in music experts and laymen: An MEG study. Journal of Cognitive Neuroscience 21, 2230–2244.

Burns, E.M., Campbell, S.L., 1994. Frequency and frequency-ratio resolution by possessors of absolute and relative pitch: examples of categorical perception. Journal of the Acoustical Society of America 96, 2704–2719.

Burns, E.M., Ward, W.D., 1978. Categorical perception - phenomenon or epiphenomenon: Evidence from experiments in the perception of melodic musical intervals. Journal of the Acoustical Society of America 63, 456–468.

Calcus, A., Lorenzi, C., Collet, G., Colin, C., Kolinsky, R., 2016. Is there a relationship between speech identification in noise and categorical perception in children with dyslexia? Journal of Speech, Language, and Hearing Research 59, 835–852.

Chang, E.F., Rieger, J.W., Johnson, K., Berger, M.S., Barbaro, N.M., Knight, R.T., 2010. Categorical speech representation in human superior temporal gyrus. Nature Neuroscience 13, 1428–1432.

Chevillet, M. A., Jiang, X., Rauschecker, J.P., Riesenhuber, M., 2013. Automatic phoneme category selectivity in the dorsal auditory stream. Journal of Neuroscience 33, 5208 –5215.

DeWitt, I., Rauschecker, J.P., 2012. Phoneme and word recognition in the auditory ventral stream. Proceedings of the National Academy of Sciences of the United States of America 109, E505–514.

Dhamala, M., Rangarajan, G., Ding, M., 2008. Analyzing information flow in brain networks with nonparametric Granger causality. Neuroimage 41, 354–362.

Di Liberto, G.M., Lalor, E.C., Millman, R.E., 2018. Causal cortical dynamics of a predictive enhancement of speech intelligibility. Neuroimage 166, 247–258.

Dick, F., Lee, H., Nusbaum, H.C., J Price, C., 2011. Auditory-motor expertise alters “speech selectivity” in professional musicians and actors. Cerebral Cortex 21, 938–948.

Du, Y., Buchsbaum, B.R., Grady, C.L., Alain, C., 2014. Noise differentially impacts phoneme representations in the auditory and speech motor systems. Proceedings of the National Academy of Sciences of the United States of America 111, 1–6.

Du, Y., Zatorre, R.J., 2017. Musical training sharpens and bonds ears and tongue to hear speech better. Proceedings of the National Academy of Sciences of the United States of America 114, 13579–13584.

Eimas, P.D., Siqueland, E.R., Jusczyk, P., Vigorito, J., 1971. Speech perception in infants. Science 171, 303–306.

Elmer, S., Meyer, M., Jancke, L., 2012. Neurofunctional and behavioral correlates of phonetic and temporal categorization in musically trained and untrained subjects. Cerebral Cortex 22, 650–658.

Feng, G., Gan, S., Wan, S., Wong, P.C.M., Chandrasekaran, B., 2018. Task-general and acousticinvariant neural representation of speech categories in the human brain. Cerebral Cortex 28, 3241–3254.

Fonov, V.S., Evans, A.C., McKinstry, R.C., Almli, C.R., Collins, D.L., 2009. Unbiased nonlinear average age-appropriate brain templates from birth to adulthood. Neuroimage 47, S102.

Freedman, D.J., Riesenhuber, M., Poggio, T., Miller, E.K., 2001. Categorical representation of visual stimuli in the primate prefrontal cortex. Science 291, 312–316.

Frey, A., François, C., Chobert, J., Velay, J.-L., Habib, M., Besson, M., 2019. Music training positively influences the preattentive perception of voice onset time in children with dyslexia: A longitudinal study. Brain Sciences 9, 91.

Geweke, J., 1982. Measurement of linear dependence and feedback between multiple time series. Journal of the American Statistical Association 77, 304–313.

Giraud, A.L., Kell, C., Thierfelder, C., Sterzer, P., Russ, M.O., Preibisch, C., Kleinschmidt, A., 2004. Contributions of sensory input, auditory search and verbal comprehension to cortical activity during speech processing. Cereb Cortex 14, 247–255.

Golestani, N., Paus, T., Zatorre, R.J., 2002. Anatomical correlates of learning novel speech sounds. Neuron 35, 997–1010.

Golestani, N., Zatorre, R.J., 2004. Learning new sounds of speech: reallocation of neural substrates. Neuroimage 21, 494–506.

Granger, C., 1969. Investigating causal relations by econometric models and cross-spectral methods. Econometrica 37, 424–438.

Guenther, F.H., Gjaja, M.N., 1996. The perceptual magnet effect as an emergent property of neural map formation. Journal of the Acoustical Society of America 100, 1111–1121.

Hakvoort, B., de Bree, E., van der Leij, A., Maassen, B., van Setten, E., Maurits, N., van Zuijen, T.L., 2016. The role of categorical speech perception and phonological processing in familial risk children with and without dyslexia. Journal of Speech, Language, and Hearing Research 59, 1448–1460.

Harnad, S.R., 1987. Psychophysical and cognitive aspects of categorical perception: A critical overview. In: Harnad, S.R. (Ed.), Categorical perception: The Groundwork of Cognition. Cambridge University Press, New York.

Herholz, S.C., Zatorre, R.J., 2012. Musical training as a framework for brain plasticity: Behavior, function, and structure. Neuron 76, 486–502.

Hickok, G., Poeppel, D., 2004. Dorsal and ventral streams: A framework for understanding aspects of the functional anatomy of language. Cognition 92, 67–99.

Howard, D., Rosen, S., Broad, V., 1992. Major/Minor triad identification and discrimination by musically trained and untrained listeners. Music Perception 10, 205–220.

Iordanov, T., Bornfleth, H., Hoechstetter, K., Lanfer, B., Scherg, M., 2016. Performance of Cortical LORETA and Cortical CLARA Applied to MEG Data. Biomag 2016.

Iordanov, T., Hoechstetter, K., Berg, P., Paul-Jordanov, I., Scherg, M., 2014. CLARA: classical LORETA analysis recursively applied. OHBM 2014.

Iverson, P., Kuhl, P.K., Akahane-Yamada, R., Diesch, E., Tohkura, Y., Kettermann, A., Siebert, C., 2003. A perceptual interference account of acquisition difficulties for non-native phonemes. Cognition 87, B47–57.

Jiang, X., Chevillet, M.A., Rauschecker, J.P., Riesenhuber, M., 2018. Training humans to categorize monkey calls: Auditory feature- and category-selective neural tuning changes. Neuron 98, 405-416.e404.

Klein, M.E., Zatorre, R.J., 2011. A role for the right superior temporal sulcus in categorical perception of musical chords. Neuropsychologia 49, 878–887.

Kraus, N., Chandrasekaran, B., 2010. Music training for the development of auditory skills. Nature Reviews Neuroscience 11, 599–605.

Kuhl, P.K., 2004. Early language acquisition: Cracking the speech code. Nature Reviews Neuroscience 5, 831–843.

Kuhl, P.K., Williams, K.A., Lacerda, F., Stevens, K.N., Lindblom, B., 1992. Linguistic experience alters phonetic perception in infants by 6 months of age. Science 255, 606–608.

Kuhnis, J., Elmer, S., Jancke, L., 2014. Auditory evoked responses in musicians during passive vowel listening are modulated by functional connectivity between bilateral auditory-related brain regions. Journal of Cognitive Neuroscience 26, 2750–2761.

Lee, Y.-S., Turkeltaub, P., Granger, R., Raizada, R.D.S., 2012. Categorical speech processing in Broca’s area: An fMRI study using multivariate pattern-based analysis. Journal of Neuroscience 32, 3942–3948.

Lehmann, D., Skrandies, W., 1980. Reference-free identification of components of checkerboard-evoked multichannel potential fields. Electroencephalography and Clinical Neurophysiology 48, 609–621.

Liebenthal, E., Desai, R., Ellingson, M.M., Ramachandran, B., Desai, A., Binder, J.R., 2010. Specialization along the left superior temporal sulcus for auditory categorization. Cerebral Cortex 20, 2958–2970.

Locke, S., Kellar, L., 1973. Categorical perception in a non-linguistic mode. Cortex 9, 355–369.

Luthra, S., Guediche, S., Blumstein, S.E., Myers, E.B., 2019. Neural substrates of subphonemic variation and lexical competition in spoken word recognition. Language, Cognition and Neuroscience 34, 151–169.

Mankel, K., Bidelman, G.M., 2018. Inherent auditory skills rather than formal music training shape the neural encoding of speech. Proceedings of the National Academy of Sciences of the United States of America 115, 13129–13134.

Mazziotta, J.C., Toga, A.W., Evans, A., Lancaster, J.L., Fox, P.T., 1995. A probabilistic atlas of the human brain: Theory and rationale for its development. Neuroimage 2, 89–101.

Messaoud-Galusi, S., Hazan, V., Rosen, S., 2011. Investigating speech perception in children with dyslexia: Is there evidence of a consistent deficit in individuals? Journal of Speech, Language, and Hearing Research 54, 1682–1701.

Meyers, E.M., Freedman, D.J., Kreiman, G., Miller, E.K., Poggio, T., 2008. Dynamic population coding of category information in inferior temporal and prefrontal cortex. Journal of Neurophysiology 100, 1407–1419.

Michel, C.M., Murray, M.M., Lantz, G., Gonzalez, S., Spinelli, L., Grave de Peralta, R., 2004. EEG source imaging. Clinical Neurophysiology 115, 2195–2222.

Miller, J.D., Wier, C.C., Pastore, R.E., Kelly, W.J., Dooling, R.J., 1976. Discrimination and labeling of noise-buzz sequences with varying noise-lead times: an example of categorical perception. Journal of the Acoustical Society of America 60, 410–417.

Moon, C., Lagercrantz, H., Kuhl, P.K., 2013. Language experienced in utero affects vowel perception after birth: A two-country study. Acta Paediatrica 102, 156–160.

Moreno, S., Bidelman, G.M., 2014. Understanding neural plasticity and cognitive benefit through the unique lens of musical training. Hearing Research 308, 84–97.

Munte, T.F., Altenmuller, E., Jancke, L., 2002. The musician’s brain as a model of neuroplasticity. Nature Reviews Neuroscience 3, 473–478.

Musacchia, G., Strait, D., Kraus, N., 2008. Relationships between behavior, brainstem and cortical encoding of seen and heard speech in musicians and non-musicians. Hearing Research 241, 34–42.

Myers, E.B., 2014. Emergence of category-level sensitivities in non-native speech sound learning. Frontiers in Neuroscience 8, 238.

Myers, E.B., Blumstein, S.E., 2008. The neural bases of the lexical effect: an fMRI investigation. Cerebral Cortex 18, 278–288.

Myers, E.B., Blumstein, S.E., Walsh, E., Eliassen, J., 2009. Inferior frontal regions underlie the perception of phonetic category invariance. Psychological Science 20, 895–903.

Noordenbos, M.W., Serniclaes, W., 2015. The categorical perception deficit in dyslexia: A meta-analysis. Scientific Studies of Reading 19, 340–359.

Oldfield, R.C., 1971. The assessment and analysis of handedness: The Edinburgh inventory. Neuropsychologia 9, 97–113.

Oostenveld, R., Fries, P., Maris, E., Schoffelen, J.M., 2011. Fieldtrip: Open source software for advanced analysis of meg, eeg, and invasive electrophysiological data. Computational Intelligence and Neuroscience 2011, 1–9.

Oostenveld, R., Praamstra, P., 2001. The five percent electrode system for high-resolution EEG and ERP measurements. Clinical Neurophysiology 112, 713–719.

Papoutsi, M., de Zwart, J.A., Jansma, J.M., Pickering, M.J., Bednar, J.A., Horwitz, B., 2009. From phonemes to articulatory codes: an fMRI study of the role of Broca’s area in speech production. Cerebral Cortex 19, 2156–2165.

Papp, N., Ktonas, P., 1977. Critical evaluation of complex demodulation techniques for the quantification of bioelectrical activity. Biomedical Sciences Instrumentation 13, 135–145.

Paraskevopoulos, E., Chalas, N., Bamidis, P.D., 2017. Functional connectivity of the cortical network supporting statistical learning in musicians and non-musicians: an MEG study. Scientific Reports 7, 16268.

Parbery-Clark, A., Skoe, E., Kraus, N., 2009. Musical experience limits the degradative effects of background noise on the neural processing of sound. Journal of Neuroscience 29, 14100–14107.

Pascual-Marqui, R.D., Esslen, M., Kochi, K., Lehmann, D., 2002. Functional imaging with low-resolution brain electromagnetic tomography (LORETA): a review. Methods and Findings in Experimental and Clinical Pharmacology 24 Suppl C, 91–95.

Phillips, C., 2001. Levels of representation in the electrophysiology of speech perception. Cognitive Science 25, 711–731.

Picton, T.W., Alain, C., Woods, D.L., John, M.S., Scherg, M., Valdes-Sosa, P., Bosch-Bayard, J., Trujillo, N.J., 1999. Intracerebral sources of human auditory-evoked potentials. Audiology and Neuro-Otology 4, 64–79.

Picton, T.W., van Roon, P., Armilio, M.L., Berg, P., Ille, N., Scherg, M., 2000. The correction of ocular artifacts: A topographic perspective. Clinical Neurophysiology 111, 53–65.

Pisoni, D.B., Luce, P.A., 1987. Acoustic-phonetic representations in word recognition. Cognition 25, 21–52.

Pisoni, D.B., Tash, J., 1974. Reaction times to comparisons within and across phonetic categories. Perception and Psychophysics 15, 285–290.

Rosen, S., Howell, P., 1987. Auditory, articulatory and learning explanations of categorical perception in speech. In: Harnad, S. (Ed.), Categorical Perception: The Groundwork of Cognition. Cambridge Univerity Press, New York.

Rozsypal, A.J., Stevenson, D.C., Hogan, J.T., 1985. Dispersion in models of categorical perception. Journal of Mathematical Psychology 29, 271–288.

Schneider, P., Scherg, M., Dosch, H.G., Specht, H.J., Gutschalk, A., Rupp, A., 2002. Morphology of Heschl’s gyrus reflects enhanced activation in the auditory cortex of musicians. Nature Neuroscience 5, 688–694.

Scott, S.K., Johnsrude, I.S., 2003. The neuroanatomical and functional organization of speech perception. Trends in Neurosciences 26, 100–107.

Seger, C.A., Miller, E.K., 2010. Category learning in the brain. Annual Review of Neuroscience 33, 203–219.

Shahin, A., Bosnyak, D.J., Trainor, L.J., Roberts, L.E., 2003. Enhancement of neuroplastic P2 and N1c auditory evoked potentials in musicians. Journal of Neuroscience 23, 5545–5552.

Shahin, A.J., Bishop, C.W., Miller, L.M., 2009. Neural mechanisms for illusory filling-in of degraded speech. Neuroimage 44, 1133–1143.

Siegel, J.A., Siegel, W., 1977. Absolute identification of notes and intervals by musicians. Perception and Psychophysics 21, 143–152.

Steinschneider, M., Fishman, Y.I., Arezzo, J.C., 2003. Representation of the voice onset time (VOT) speech parameter in population responses within primary auditory cortex of the awake monkey. Journal of the Acoustical Society of America 114, 307–321.

Steinschneider, M., Volkov, I.O., Noh, M.D., Garell, P.C., Howard, M.A., 3rd, 1999. Temporal encoding of the voice onset time phonetic parameter by field potentials recorded directly from human auditory cortex. Journal of Neurophysiology 82, 2346–2357.

Suga, N., 1989. Principles of auditory information-processing derived from neuroethology. Journal of Experimental Biology 146, 277–286.

Toscano, J.C., Anderson, N.D., Fabiani, M., Gratton, G., Garnsey, S.M., 2018. The time-course of cortical responses to speech revealed by fast optical imaging. Brain and Language 184, 32–42.

Weiss, M.W., Bidelman, G.M., 2015. Listening to the brainstem: Musicianship enhances intelligibility of subcortical representations for speech. Journal of Neuroscience 35, 1687–1691.

Wong, P.C., Skoe, E., Russo, N.M., Dees, T., Kraus, N., 2007. Musical experience shapes human brainstem encoding of linguistic pitch patterns. Nature Neuroscience 10, 420–422.

Xu, Y., Gandour, J.T., Francis, A., 2006. Effects of language experience and stimulus complexity on the categorical perception of pitch direction. Journal of the Acoustical Society of America 120, 1063–1074.

Yi, H.G., Chandrasekaran, B., 2016. Auditory categories with separable decision boundaries are learned faster with full feedback than with minimal feedback. Journal of the Acoustical Society of America 140, 1332–1332.

Yi, H.G., Maddox, W.T., Mumford, J.A., Chandrasekaran, B., 2016. The role of corticostriatal systems in speech category learning. Cerebral Cortex 26, 1409–1420.

Yoo, J., Bidelman, G.M., 2019. Linguistic, perceptual, and cognitive factors underlying musicians’ benefits in noise-degraded speech perception. Hearing Research 377, 189–195.

Zatorre, R., Halpern, A.R., 1979. Identification, discrimination, and selective adaptation of simultaneous musical intervals. Perception and Psychophysics 26, 384–395.

Zatorre, R., McGill, J., 2005. Music, the food of neuroscience? Nature 434, 312–315.

Zatorre, R.J., 1983. Category-boundary effects and speeded sorting with a harmonic musical-interval continuum: Evidence for dual processing. Journal of Experimental Psychology: Human Perception and Performance 9, 739–752.

Zendel, B.R., Alain, C., 2009. Concurrent sound segregation is enhanced in musicians. Journal of Cognitive Neuroscience 21, 1488–1498.

Zoubrinetzky, R., Collet, G., Serniclaes, W., Nguyen-Morel, M.-A., Valdois, S., 2016. Relationships between categorical perception of phonemes, phoneme awareness, and visual attention span in developmental dyslexia. PLoS One 11, e0151015.

